# Marmoset Brain Mapping V5: an Anatomical and Connectivity Atlas of the Cerebellum

**DOI:** 10.1101/2023.02.02.526911

**Authors:** Xiaojia Zhu, Haotian Yan, Yafeng Zhan, Furui Feng, Chuanyao Wei, Yong-Gang Yao, Cirong Liu

## Abstract

The cerebellum is a key region of the brain for motor control and cognitive functioning, and it is involved bidirectional communication with the cerebral cortex. As one of the smallest non-human primates, the common marmoset provides many advantages in the study of the anatomy and the functions of cerebello-cerebral circuits. However, the cerebellum of the marmoset is far from being well described in published resources. In this study, we present a comprehensive atlas of the marmoset cerebellum in three parts: 1) fine-detailed anatomical atlases and surface-analyzing tools of the cerebellar cortex, based on ultra-high resolution *ex-vivo* MRI; 2) functional-connectivity gradients of the cerebellar cortex, based on awake resting-state fMRI; and 3) structural-connectivity based mapping of the cerebellar nuclei, based on high-resolution diffusion tractography. The atlas shows the anatomical details of the marmoset cerebellum; reveals distinct gradient patterns of the intra-cerebellar and the cerebello-cerebral functional connectivity; and maps the topological relationship of the cerebellar nuclei in cerebello-cerebral circuits. As version 5 of the *Marmoset Brain Mapping* project, the atlas describes the anatomical and connectivity details of the marmoset cerebellum and is publicly available via *marmosetbrainmapping.org*.

## Introduction

The common marmoset has risen quickly as a promising non-human primate (NHP) model for neuroscience research ^1–3^. Due to its small brain size, the marmoset provides advantages in the study of brain anatomy and the functional mapping of brain architecture. All of which has enabled the development of valuable open resources and research tools ^4–11^. Specifically, via the *Marmoset Brain Mapping* project (*marmosetbrainmapping*.*org*), we have provided ultra-high-resolution *ex-vivo* diffusion MRI and awake resting-state fMRI data of the marmoset brain ^12–15^. Based on the resource, we have produced an anatomical and functional atlas of the marmoset cerebral cortex in fine detail and created useful templates for surface-based visualization and analysis. Together with the *Marmoset Brain Connectivity* project ^15–17^, a neuronal tracing database of the marmoset neocortex, our resources greatly facilitate the study of the neuroanatomy and the connectome of the marmoset brain ^18^.

Despite the recent progress and the public availability of the resources, one of the major brain regions, the cerebellum, has only be described in histological atlas books, including the Paxinos atlas ^19^ and the Hardman atlas ^20^, but not been mapped in the earlier published digital atlases for imaging and connectome studies. The cerebellum, historically regarded as a motor-control region, is now found to be a critical node in neural circuits involved in a wide range of cognitive functions ^21–24^. These functions involve bidirectional communication between the neocortex and the cerebellum ^24^. In particular, recent studies used the diffusion embedding, a non-linear dimensional reduction method, to obtain the gradients in the similarity matrices of resting-state fMRI functional connectivity (functional gradient) of the cerebellar cortex ^25,26^. These gradients reflected spatial distribution patterns of the variances in the functional connectivity of the cerebellar cortex and were associated with distinct brain networks and diverse task domains ^22,25,27^. Although the cerebellar cortex has a remarkably uniform cytoarchitecture and stereotypic local circuitry, it shows a rich diversity of region-specific connectivity to the neocortex, the description of which is essential to the understanding of the functional dynamics and the neural circuits that underlie motor-control and cognitive functioning ^28,29^.

In the current study, we have aimed to develop a comprehensive atlas of the marmoset cerebellum by using the multi-modal MRI data of the *Marmoset Brain Mapping* project (*marmosetbrainmapping*.*org*). First, based on our ultra-high resolution *ex-vivo* MRI, we developed an anatomical atlas of the cerebellar cortex and high-resolution surfaces that preserved fine-grained folia patterns. Second, using our recently released resources of awake resting-state fMRI ^15^, we examined the functional connectivity patterns of the cerebellum and compared the gradient patterns of the intra-cerebellar and the cerebello-cerebral functional connectivity. Last, based on diffusion tractography and functional connectivity analysis, we performed connectivity-based mapping of the cerebellar nuclei, which reflected nucleus-specific connectivity patterns in cerebellar-cerebral circuits. The cerebellar atlases can be downloaded for data analysis or interactively viewed online, utilizing the web-based atlas viewer of the Marmoset Brain Mapping Project. The study greatly extends the available neuroimaging resources of the marmoset brain and will accelerate comparative studies of the cerebellum.

## Results

### Fine-detailed surface reconstruction and lobular parcellations of the cerebellar cortex

We constructed a fine-grained surface and lobular parcellation of the marmoset cerebellar cortex based on ultra-high resolution *ex-vivo* MRI data ^13^. Although the marmoset has an almost smooth neocortex, its cerebellar cortex is tightly folded into a complex folia pattern (**Fig. 1**). As the cerebellar cortex is a thin and folded sheet of neuronal tissue, a folium of the marmoset is hard to resolve by normal *in-vivo* MRI due to the partial volume effects (**Fig. 1A-B**). Thus, *ex-vivo* MRI with ultra-high resolution becomes essential to describe the detailed 3D folding patterns of the cerebellar cortex. As shown in **Fig. 1**, our ultra-high resolution MRI data ^13^ were able to resolve the detailed pattern of an individual folium and the structure of its layers.

**Figure 1.**
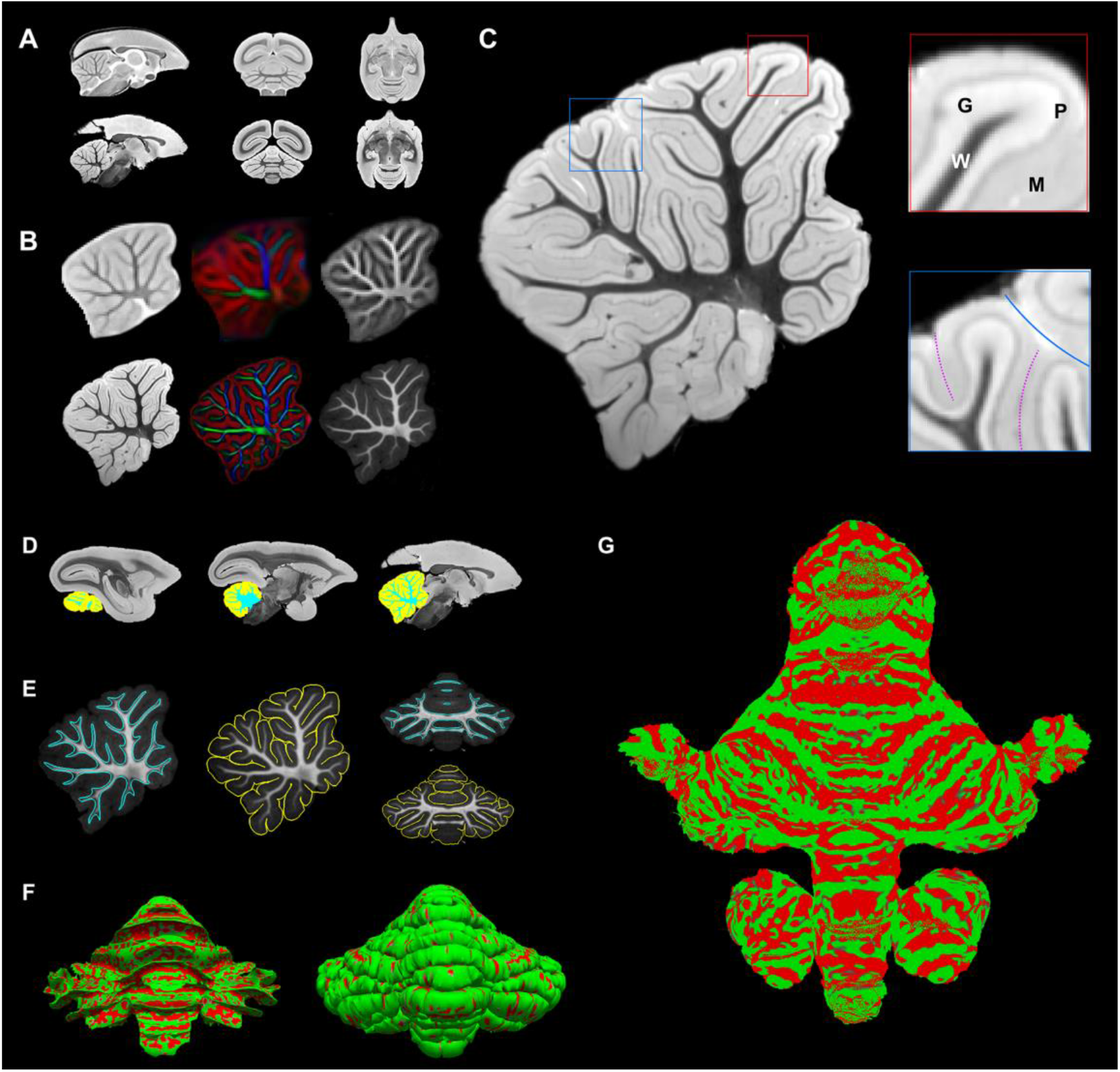
Ultra-high resolution *ex-vivo* MRI reveals anatomical details of the marmoset cerebellum. **(A)** The top is the *in-vivo* 200 μm T2-weighted population template; and the bottom is the *ex-vivo* 50 μm T2^*^-weighted image. (**B**) The middle-sagittal slice of the cerebellum. The top shows the T2-weighted, diffusion tensor-based directionally encoded color (DEC), and the T1-weighted images (from left to right) of the *in-vivo* 200 μm images. The bottom displays the T2^*^-weighted, DEC image, and the magnetization transfer ratio (MTR; T1-weight-like contrast) of the *ex-vivo* 80 μm images. (**C**) The zoom-in view of a middle-sagittal slice of the cerebellum. Our ultra-high resolution *ex-vivo* image reveals the layer pattern (an example highlighted in the red box) and the folia pattern (highlighted in the blue box). W: white matter; G: Granular layer; P: Purkinje layer M: Molecular layer. Blue solid line: big lobular boundary; purple dash lines: boundaries of small folia. (**D**) The segmentation of the cerebellum into white matter (blue) and gray matter (yellow). From left to right shows sagittal slices of the T2^*^-weighted image. (**E**) The surface outlines are displayed on the *ex-vivo* MTR image. A middle sagittal slice and a coronal slice show the white matter surface outline in blue and the pial surface in yellow. (**F**) The 3D view of the white matter surface (left) and the pial surface (right). The curvature of the cortex is displayed as the overlay. (**G**) Flat map of the cerebellar cortex, generated from the surface displayed in **E**. Colored overlay shows the curvature of the cortex, with gyri in green and sulci in red.

Based on the ultra-high resolution data, we manually segmented the cerebellum into gray matter and white matter (**Fig. 1D**), and then reconstructed the folia-level pial and white matter surfaces using Freesurfer ^30^ (**Fig. 1E-F**). As the pial surface was expanded from the white matter surface, the vertices were paired and matched on the two surfaces. To show the hierarchical levels of folding patterns from large lobules to small folia, the surfaces were densely tessellated with a total of 286308 vertices and 572724 faces. As the detailed folding patterns complicated the visualization of the cerebellar data, we unfolded the pial surface and created the flat maps of the marmoset cerebellar cortex (**Fig. 1G**). The flat map was built on dense-tessellated surfaces, which were unfolded down to the level of individual folium. The dense-tessellated flat map fully took advantage of the high-resolution MRI data to preserve the anatomical details of the cerebellar cortex.

We manually delineated the cerebellar lobules on the ultra-high resolution templates according to the nomenclature of the Paxinos atlas ^19^. The cerebellar cortex was parcellated into the 17 lobules: Lingula I (I), Central lobule II (II), Culmen III (III), Declive IV (IV), lobule V (V), Folium VI (VI), Tuber VII (VII), Pyramid VIII (VIII), Uvula IX (IX), Nodulus X (X), Simplex lobule (SIM), Paramedian lobule (Par), Copula (Cop), Flocculus (Fl), Paraflocculus (PFI), Crus I and Crus II (**Fig. 2A**). For convention, we merged lobules I, II, III, and IV into one region (I-IV) and IX and X into the vestibulocerebellar region (IX-X) to form a 13-lobule parcellation (**Fig. 2A-C**), following a similar strategy as described in a study for the human cerebellar atlas ^31^. The anatomical lobular atlas was mapped onto the surfaces and the flat map to facilitate surface-based analysis (**Fig. 2D**).

**Figure 2.**
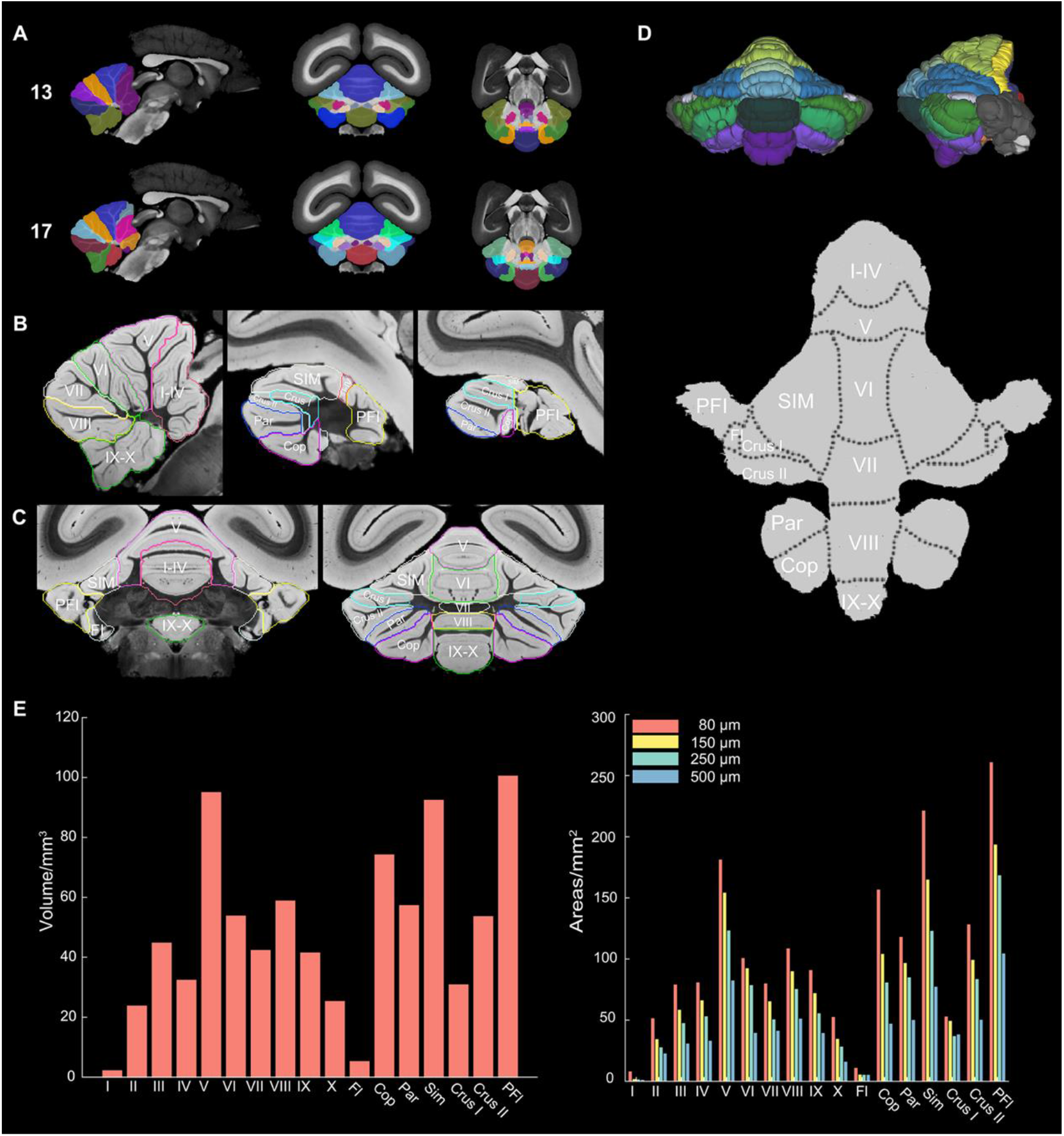
The lobular parcellation of the cerebellar cortex. **(A)** The atlas of lobular parcellation. The top is the 13-lobular parcellation and the bottom is the 17-lobular parcellation. From left and right are examples of a sagittal, a coronal, and a horizontal slice. The underlying image is an *ex-vivo* MTR image. **(B)** Examples of three sagittal slices of the 13-lobular parcellation on an *ex-vivo* T2^*^-weighted image. (**C**) Examples of two coronal slices. The 13-lobules include I-IV, V, VI, VII, VIII, IX-X, SIM, Par, Cop, Fl, PFI, Crus I and Crus II. (**D**) The 13-lobular parcellation on the surface (upper panel) and flat map of the cerebral cortex (below). (**E**) The volume size of the 17 cerebellar lobules (left) and the estimated surface areas using data with different resolutions (right). Detailed values of sizes are provided in **Table 1** and **Supplementary Table S1**.

The high-resolution volumetric and surface atlases allowed us to accurately estimate the volume and surface area of the whole cerebellar cortex and each lobule. After applying a brain-shrinkage correction (see Methods), the total volume and area of the cerebellar cortex are 838.3 mm^3^ and 1782.8 mm^2^ at an 80 μm isotropic resolution, respectively (**Fig. 2E** and **Table 1**). Note that we also estimated the surface area at a higher resolution (a 50 μm isotropic T2* structural image), and obtained a surface area of 1812.7 mm^2^, close to the estimation based on 80 μm multi-modal data. The result suggests that the 80 μm resolution was sufficient to distinguish most individual folia. The three largest lobules are PFl, SIM, and lobule V, and the three smallest lobules are lobule I, Fl, and lobule II (Note: the flat map did not reflect the surface area of the lobules due to the surface flattening). We also measured the surface area of each lobule on the downsampled data (**Fig. 2E**, right). For the lower resolution in which the folia failed to be fully reconstructed, the surface area was largely underestimated. For example, we downsampled our data into 150 μm isotropic, 250 μm (typical *in-vivo* structural data of the marmoset brain), and 500 μm (typical *in-vivo* diffusion MRI data), and their estimated surface areas were reduced to 1383.0 mm^2^ (77.6% of the original size), 1124.2 mm^2^ (63.1%), 730.4 mm^2^ (41.0%), respectively (**Fig. 2E** and **Supplementary Table S1**). Thus, the high resolution was essential to reveal the folia pattern and estimate the accurate area size of the cerebellar cortex ^32^.

**Table 1.** The volume and surface area of 17 lobules.

| Lobules | I | II | III | IV | V | VI | VII | VIII | IX |
| --- | --- | --- | --- | --- | --- | --- | --- | --- | --- |
| Volume/mm<sup>3</sup> | 2.44 | 24.03 | 45.04 | 32.63 | 95.27 | 54.06 | 42.57 | 59.07 | 41.74 |
| Area/mm<sup>2</sup> | 8.04 | 51.49 | 79.09 | 80.88 | 181.25 | 100.72 | 79.98 | 108.58 | 90.94 |
| Lobules | X | SIM | Par | Cop | FI | PFI | Crus I | Crus II | Total |
| Volume/mm<sup>3</sup> | 25.52 | 92.66 | 57.53 | 74.45 | 5.49 | 100.76 | 31.10 | 53.84 | 838.30 |
| Area/mm<sup>2</sup> | 52.58 | 221.35 | 118.06 | 156.76 | 10.97 | 260.86 | 52.92 | 128.26 | 1782.80 |

Note: The 17 lobules in marmoset cerebellar cortex are parcellated following the nomenclature of the Paxinos atlas <sup>19</sup> and the volumes are estimated from ultra-high resolution *ev-vivo* data of one marmoset <sup>13</sup>.

### Distinct functional gradients of the intra-cerebellar connectivity and the cerebello-cerebral connectivity

Previous human studies have revealed that the cerebellar cortex shows distinct topographic patterns of intrinsic functional connectivity more relevant to its functional domains than to the anatomical lobular parcellations ^22,25^. To reveal the functional connectivity pattern of the marmoset cerebellar cortex, we calculated the functional gradients by analyzing the intra-cerebellar functional connectivity and the cerebello-cerebral connectivity using diffusion map embedding ^27,33,34^.

We first examined the principal gradients (gradient-1), which accounted for the largest part of the variability in the resting-state connectivity patterns (**Supplementary Fig. S1A**). For gradient-1 of the intra-cerebellar functional connectivity (**Fig. 3A**), the negative extreme (blue) was in areas of motor representation (lobules I-VI) and gradually extended to areas of non-motor representation with the positive extreme (orange) located in Crus I-II and Par. The gradient-1 of the cerebello-cerebral connectivity was more complex and had one extreme in cerebellar vermis and lobules I-VI, which extended to Crus I and Crus II and reached the Sim and PFI as the other extreme. Other gradients of the intra-cerebellar and cerebello-cerebral functional connectivity also showed different patterns in different lobules (**Fig. 3A**). For example, gradients 2 and 3 of the intra-cerebellar connectivity were asymmetrical, with their positive extremes located in the lobules VI and the Crus I-II, respectively. On the contrary, gradients 2 and 3 of the cerebello-cerebral connectivity were symmetrical with more complex gradient-changing patterns as the gradient-1. We quantitatively analyzed the spatial relationship between the intra-cerebellar and the cerebello-cerebral gradients in both marmosets and humans. The analysis revealed differences between the two species in terms of the similarity of these gradients (**Fig. 3B**). In humans, the first four gradients of both types demonstrated high similarity, with the first two gradients being nearly identical (with R values exceeding 0.9), indicating a strong influence of cerebello-cerebral connections on intra-cerebellar connectivity ^25^. In contrast, the results for marmosets exhibited weaker relationships between the two types of gradients. The percentage of data variance explained by the two types of gradients was also similar in humans but not in marmosets (**Supplementary Fig. S1**). Furthermore, in marmosets, the fourth intra-cerebellar gradient (4.75% of variance) demonstrated weak similarity (with an R value of 0.62) to the first cerebello-cerebral gradient (35.19% of variance). The third intra-cerebellar gradient (5.57% of variance) displayed weak similarity (with an R value of 0.6) to the fourth cerebello-cerebral gradient (3.11% of variance). While these similarities in marmoset gradient patterns are weaker than those observed in humans, they suggest that the influence of cerebello-cerebral connections on the intrinsic functional connectivity of the cerebellar cortex also exists in marmosets, albeit to a much lesser extent. These results highlight the evolutionary divergence in the functional connectivity between the neocortex and cerebellar cortex in the two primate species examined.

**Figure 3.**
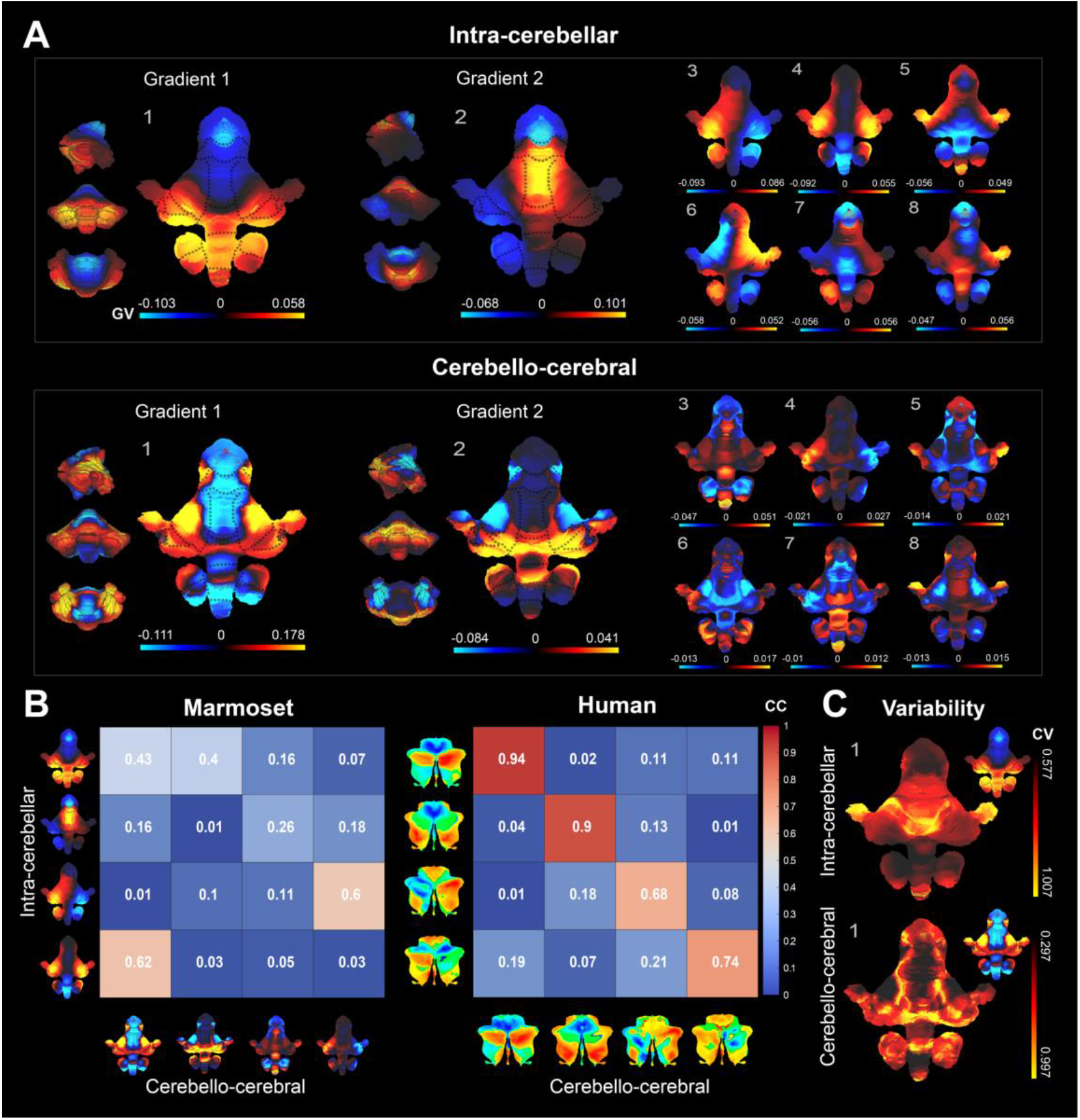
Functional gradients of the intra-cerebellar and the cerebello-cerebral connectivity. (**A**) Gradient-1 to 8 of the two types of connectivity. (**B**) The corresponding gradients of the two types show similar spatial patterns in humans (right) but not in marmosets (left). The numbers and color are correlation coefficients (CC) between the spatial patterns of two gradients. (**C**) The individual variability of the intra-cerebellar gradient-1 (top) and the cerebello-cerebral gradient-1 (bottom); The color bar represents coefficients of variability (CV). The gradient itself is also displayed on the right corner of each panel. The variabilities of other gradients (2 to 8) are displayed in the **Supplementary Fig. S2**.

Due to the large individual variability in the functional connectivity of marmoset brains, the population-level functional gradients may not capture the individual functional connectivity patterns of the cerebellum. Based on awake resting-state fMRI of 39 marmosets, we examined the individual variability of both the intra-cerebellar and the cerebello-cerebral principal gradients (**Fig. 3C** and **Supplementary Fig. S2**). For both types of gradients, their two extremes had the lowest individual-level variability (the darkest areas), while the areas with the highest variabilities were mainly located at the boundary of two extremes. The result indicated that the functional connectivity patterns in the gradient extremes were highly consistent across individuals and the variability increased gradually from two extremes to other areas.

To characterize the spatial distribution of the gradient extremes, we mapped the top 10% values of gradient-1, the top 10% values of gradient-2, and the lowest 10% values of gradient-1 (**Fig. 4**, left) on lobular parcellations. For the intra-cerebellar gradients (**Fig. 4A**), the top 10% values were exclusively located at the non-motor lobules, including VII-X, Fl, PFl, Cop, PM, Sim, Crus I, and Crus II; and the lowest 10% values were in motor areas, including lobules I-VI. The patterns were also reflected by spectral clustering on intra-cerebellar functional connectivity (**Supplementary Fig. S3**). In other words, the two extremes of the intra-cerebellar gradients were exclusively located in either non-motor or motor areas. On the contrary, the top values of the cerebello-cerebral gradients were in both the motor and non-motor areas (**Fig. 4B**), further indicating the differences between the two types of gradients.

**Figure 4.**
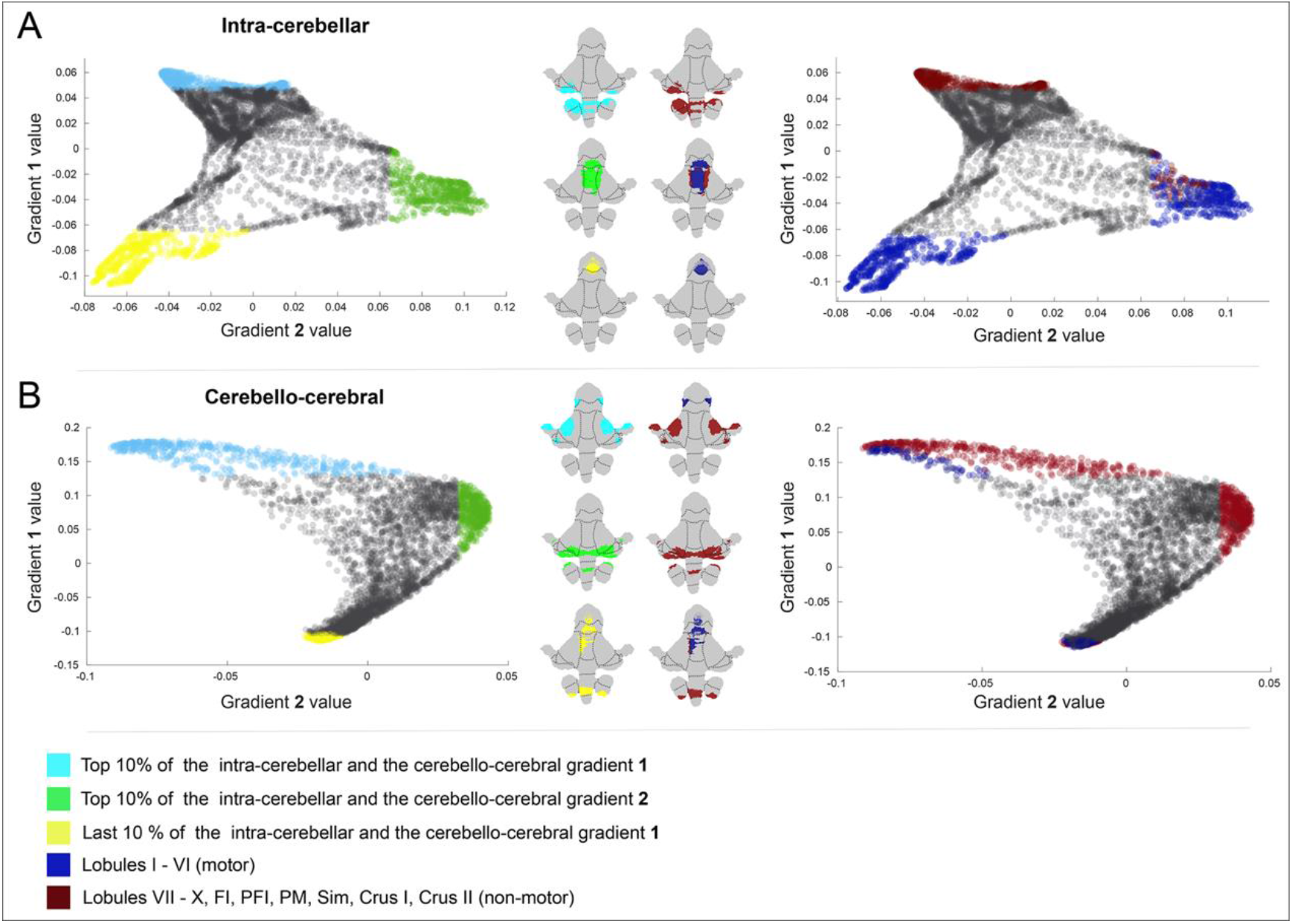
The spatial distributions of gradient extremes. (**A-B**) Scatter maps and regional distributions of the gradient extremes in the first two intra-cerebellar functional gradients (**A**) and the cerebello-cerebral functional gradients (**B**).

The above gradient analyses showed the topological patterns of the cerebello-cerebral functional connectivity but did not demonstrate which cerebral regions connected with the cerebellum. In our previous studies, we provided a comprehensive mapping of the functional networks of the marmoset cerebral cortex ^15^. Here, we calculated the functional connectivity of each cerebellar vertex to these cerebral networks to obtain the cerebellar connectivity maps of each network (**Fig. 5** and **Supplementary Fig. S4**). Interestingly, the top extremes of the cerebello-cerebral gradient-1 shown in **Fig. 4B** were also the cerebellar areas that had the highest connectivity to most cerebral networks (**Fig. 5A**), highlighting the importance of these areas for cerebello-cerebral communications. To verify the pattern, we calculated the variation of correlations to all fifteen networks that were defined in our recent study ^15^ for each cerebellar vertex and obtained a map of variation (**Fig. 5B**), where low values represented consistent connectivity across different networks and *vice versa*. The top extremes of the cerebello-cerebral gradient-1 demonstrated the relatively strong and robust functional connectivity to most of the cortical networks than the other areas according to the variation map. The map showed a similar spatial pattern to the gradient-1 of the cerebello-cerebral connectivity but not to that of the intra-cerebellar connectivity, indicating the divergence of the two types of functional connectivity of the cerebellar cortex.

**Figure 5.**
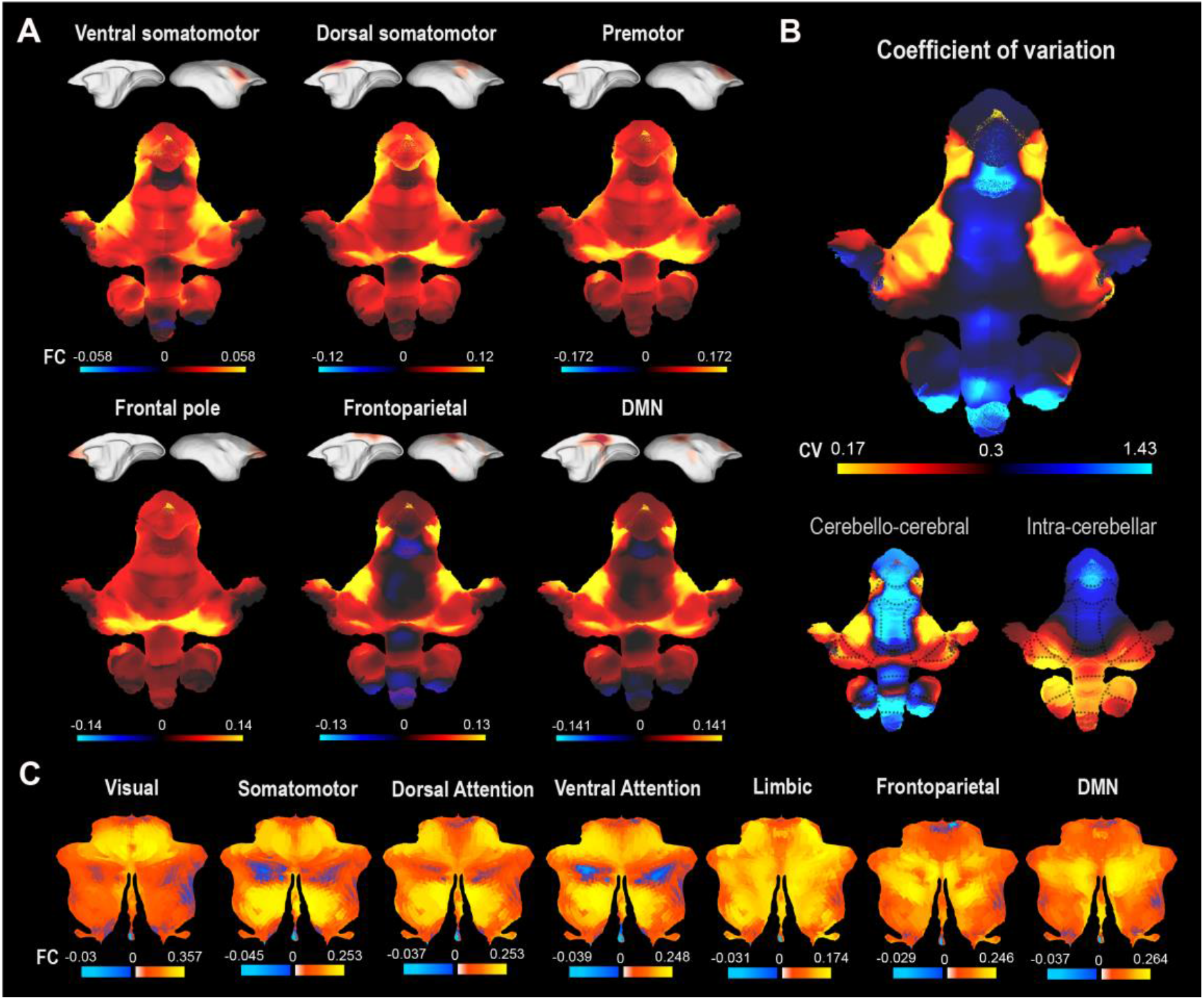
Functional connectivity patterns between the cerebral functional networks and the cerebellar cortex. (**A**) The functional connectivity between the cerebellar cortex and six selected cerebral networks. The connectivity maps to all cerebral networks are displayed in the **Supplementary Fig. S4**. The cerebral networks were defined by our previous study ^15^. DMN: default model network. (**B**) The variation map of the connectivity strengths to the cerebellar cortex across all cerebral networks (top). The map shows a similar spatial pattern as the cerebello-cerebral functional gradient-1 (bottom-left) but differs to the intra-cerebellar functional gradients-1 (bottom-right). (**C**) The functional connectivity between the cerebellar cortex and the seven cerebral networks ^35^ in humans, shown on the flat map of human cerebellar cortex ^31^.

Interestingly, in marmosets, the frontoparietal network and the default mode network exhibit a significant overlap in terms of cerebellar connectivity (R = 0.98), despite being spatially separated in the cortex. We further analyzed the functional connectivity between the cerebellar cortex and the seven cerebral networks of humans ^35^ (**Fig.5C**). In comparison to marmosets, the frontoparietal network and default mode network in humans had a lower similarity, although they remained similar (R = 0.70), suggesting that the two networks may have diverged during the primate evolution and become more distinguishable to the cerebellum in humans. Additionally, in comparison to marmosets, the functional connectivity between the cerebellum and the cerebral cortex was stronger in humans, supporting the notion that the intra-cerebellar gradients and cerebello-cerebral gradients are more similar in humans due to stronger connections between the cerebral cortex and cerebellar cortex.

### Structural-connectivity-based mapping reveals two extremes of the cerebellar nuclei in cerebello-cerebral circuits

In addition to the cerebellar cortex, cerebellar nuclei are the other substructures of the cerebellum. They are, from lateral to medial: the dentate nucleus, the interposed nucleus, and the fastigial nucleus. To characterize these nuclei, we constructed a 3D atlas on our high-resolution *ex-vivo* templates and estimated their volumes (**Fig. 6A-B**). The dentate nucleus was the largest and its volume (13.72 mm^3^) was more than the other two nuclei combined (interposed nucleus:5.53 mm^3^; fastigial nucleus: 5.28 mm^3^). The size ratio of the dentate nucleus in the marmoset was higher than in the mouse but was similar to that of the rhesus monkey (**Fig. 6C**), demonstrating the enlargement of the dentate nucleus in primate brains.

**Figure 6.**
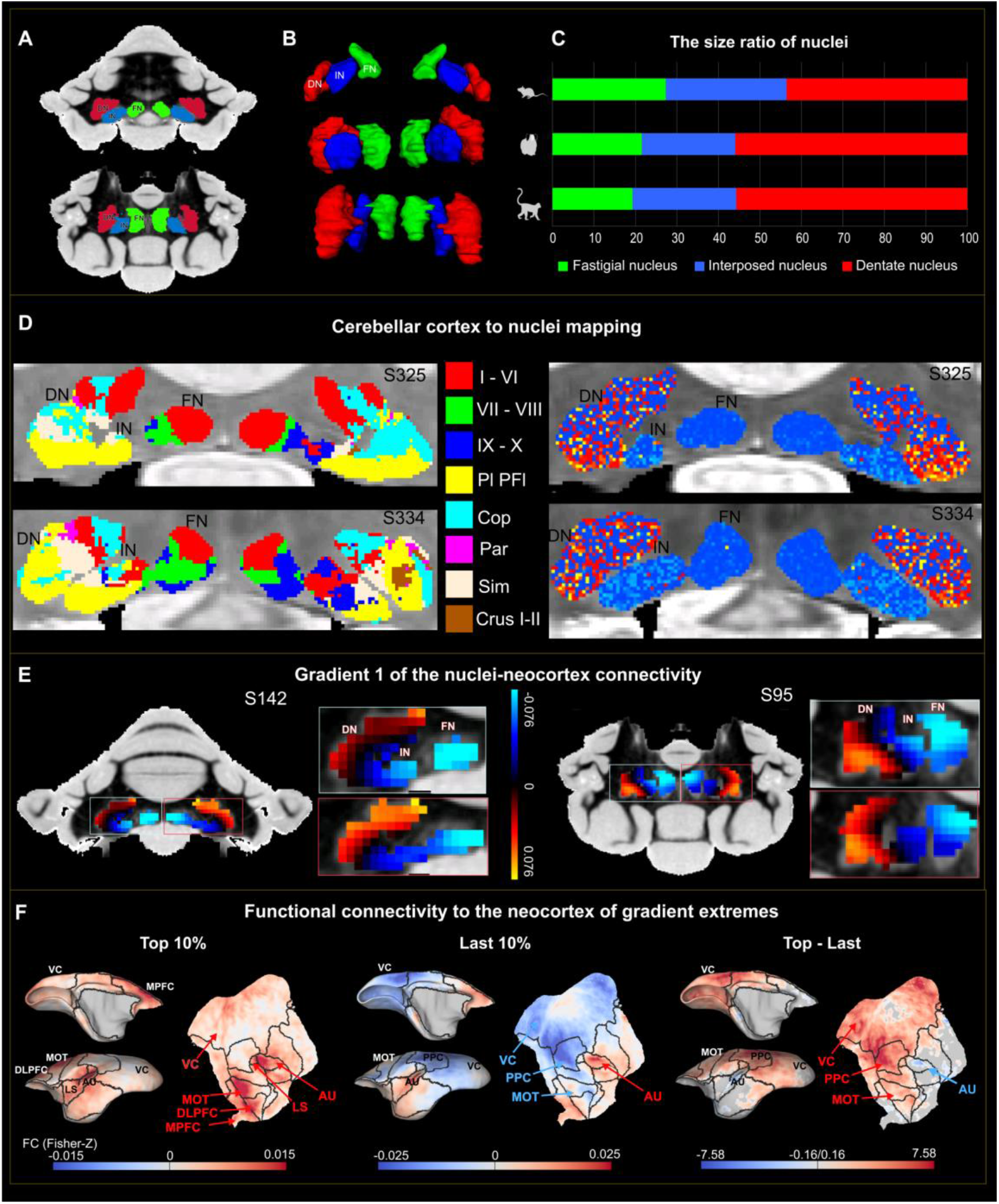
The structural-connectivity-based mapping and functional gradients of the cerebellar nuclei. (**A**) The anatomical atlas of the cerebellar nuclei of the marmoset. The top is a representative coronal slice, and the bottom is a horizontal slice. DN: dentate nucleus (red); IN: interposed nucleus (blue); FN: fastigial nucleus (green). (**B**) The 3D view of the three cerebellar nuclei of the mouse (top), the marmoset (middle), and the macaque (bottom). (**C**) The relative size ratio (in percentages) of three nuclei in the three species (top, mouse; middle, marmoset; bottom, macaque). (**D**) Mapping the properties of the cerebellar cortex to the nuclei based on their structural connectivity using 80 μm isotropic data. The left shows the cerebellar-lobule labels mapped, and the right is the cerebello-cerebral functional gradient-1 mapped on the nuclei. For each panel, the top is the coronal slice 325 and the bottom is the slice 334. More slices are shown in the **Supplementary Fig.S5**. (**E**) Gradient-1 of the functional connectivity between the cerebellar nuclei and the cerebral cortex (nuclei-neocortex gradient-1). For each zoom-in panel, the top shows the nuclei-neocortex gradient-1 of the left nuclei (blue box), and the bottom is the right nuclei (red box). (**F**) Functional connectivity (Fisher Z-converted) patterns of two extremes (top 10 % and last 10% value of the nuclei-neocortex gradient-1) to the left cerebral cortex. The statistical difference was evaluated by paired two-sample t-test (p<0.05, FDR corrected). The results to the right cerebral cortex are shown in the **Supplementary Fig. S7**. MOT: motor and premotor cortical regions, DLPFC: dorsolateral prefrontal cortex, MPFC: medial prefrontal cortex, LS: insular cortex, and AU: auditory cortex.

The cerebellar nuclei route information from different parts of the cerebellar cortex to other brain regions. Little is known for the connection between the cerebellar nuclei and the cerebellar cortex in the marmoset brain. Based on diffusion tractography, we estimated the structural connectivity between the nuclei and the cerebellar cortex to obtain their connectivity profiles. We then used the profiles to associate each nucleus voxel with the most-connected voxel of the cerebellar cortex (**Fig. 6D**). According to the relationship, we assigned each nucleus voxel with the properties of its associated cerebellar-cortical voxel, including the value of the gradient-1 of the cerebello-cerebral functional connectivity and the labels of the lobule parcellation. The structural-connectivity-based mapping was performed on two *ex-vivo* data, one is the 80 μm isotropic (**Fig. 6D**), and the other 150 μm isotropic (**Supplementary Fig. S6**). Both sets of data produced consistent results, but a higher spatial resolution (80 μm) generated a more fine-grained mapping.

Most voxels of the dentate nucleus were assigned with one extreme (high values) of the cerebello-cerebral functional gradients, and mainly stemmed from non-motor areas of the cerebellar cortex (**Fig. 6D**). On the contrary, the fastigial nucleus occupied the other extreme (low values) of the cerebello-cerebral gradients. The assignment of cerebellar lobules also confirmed the connectivity patterns between the cerebellar cortex and nuclei. The fastigial nucleus was only connected to the vermis regions (lobules I-X) of the cerebellar cortex for motor functions, while the interposed nucleus and the dentate nucleus connect to more hemispheric areas such as Sim and Crus I and Crus II for non-motor functions. This observation was consistent with the result seen in the mouse using anterograde virus tracing ^36^. Overall, the structural-connectivity-based mapping revealed the topological relationship of the connections between the cerebellar cortex and the nuclei.

We then studied the topological relationship between the cerebellar nuclei and the cerebral cortex. As the projections from the cerebellar nuclei to the cerebral cortex are transsynaptic, diffusion tractography may not be able to reconstruct these indirect connections. Thus, we used the resting-state fMRI to map the cerebellar-nuclei-cerebral connections. We calculated the functional connectivity between the cerebellar nuclei and the cerebral cortex and analyzed their functional gradients. The principal gradient shows a clear continuous change from the dentate nucleus (highest extreme) to the fastigial nucleus (lowest extreme; **Fig. 6E**), similar to the patterns in the structural-connectivity mapping of the cerebellar cortex (**Fig. 6D**). To compare the functional connectivity patterns of the two extremes, we extracted the voxels with the top and the last 10% of the gradient and calculated their correlation strength to the cerebral cortex (**Fig. 6F** and **Supplementary Fig. 7**). The positive extreme (top 10%; mainly in the dentate nucleus) exhibited positive functional connectivity with most regions in the cerebral cortex, particularly in the motor and premotor cortical regions (MOT), dorsolateral prefrontal cortex (DLPFC), medial prefrontal cortex (MPFC), insular cortex (LS), and auditory cortex (AU). In contrast, the negative extreme (bottom 10%; mainly in the fastigial nucleus) displayed more negative functional connectivity with the cerebral cortex, such as the visual cortex (VC), posterior parietal cortex (PPC), and MOT. A paired two-sample t-test further confirmed the stronger functional connectivity of the dentate nucleus compared to that of the fastigial nucleus for most regions of the cerebral cortex, including the VC, PPC, and MOT. The opposite connectivity patterns of the dentate nucleus and the fastigial nucleus indicated their distinct roles in the cerebello-cerebral circuit loop and cerebellar functions.

## Discussion

In the study, we developed a comprehensive atlas of the marmoset cerebellum based on ultra-high resolution structural MRI data and awake resting-state fMRI data. This study has not only provided useful tools to facilitate neuroimaging and comparative studies of the primate cerebellum, but also revealed anatomical and functional details of the marmoset cerebellum that were previously far from being well described.

The cerebellum and the neocortex co-evolved phylogenetically and expanded significantly in primate brains ^37–39^. A previous study reported that the surface area of the human cerebellar cortex was equal to almost 80% of the total surface area of the neocortex, but in the macaque monkey was approximately 33% of its neocortex ^32^, which suggested that the cerebellum may play a prominent role in the evolution of human behaviors and cognition. Here, we measured the surface areas of the marmoset brain and found that the area of the cerebellar cortex was about 61.5% of that of the neocortex, which is lower than the ratio of humans (80%) but much higher than of the macaque (33%). These values suggested that a few species may not determine the co-evolutionary patterns between the cerebellum and the cerebral cortex, and the size ratio of which may not be a simple linear relationship during evolution. Large-scale comparative neuroimaging across diverse species becomes essential to reveal the principles of cerebellar evolution and to examine the potential functional divergence ^40,42,44,46^.

The gradient analysis can characterize functional connectivity patterns and reflect the anatomical and functional organization of the brain. The brain areas are not randomly organized but followed a center topographical organization. The topography, which shows roughly sensorimotor to trans-modal hierarchical changes, has been reported in brain cytoarchitecture, connectivity, and gene expression pattern ^33,41,43,45,47^. The diffusion embedding method has proven useful for revealing the topographical pattern by recovering a low-dimensional spatial representation (functional gradients) from high-dimensional functional connectivity data ^27,33,34,48^. Based on the diffusion embedding, our results and previous studies ^25,26^ revealed a similar functional gradient for the intra-cerebellar and the cerebello-cerebral functional connectivity of the human brain. The observation in humans suggested that the cerebello-cerebral connections may influence the intrinsic functional connectivity of cerebellar regions, which contributed to similar spatial distribution between the two types of gradients. In our study, we observed different gradient patterns of the intra-cerebellar and the cerebello-cerebral connectivity in the marmoset. Although the intra-cerebellar and the cerebello-cerebral principal gradients showed different patterns, both shared a similar topological spatial change with one extreme in areas related to motor functions and the other in the non-motor. The cerebello-cerebral principle gradient pattern also matched with the connectivity patterns of cerebral functional networks to the cerebellar cortex. These patterns demonstrated the influence of the cerebello-cerebral connections on shaping the functional connectivity pattern of the marmoset cerebellar cortex. However, these influences in the marmoset cerebellum may be not as strong as in humans, as we observed more complex gradient patterns of the cerebello-cerebral connectivity than that of the intra-cerebellar connectivity. The species difference indicates the cerebellum and its connectivity with the cerebral cortex underwent prominent changes during primate evolution.

The cerebello-cerebral gradient reflected the functional connectivity patterns of different cerebellar networks to the cerebellar cortex in marmosets. One interesting observation was that the functional connectivity patterns of the frontoparietal-like and default-mode-like networks appeared to highly similar in the marmoset cerebellum (**Figure 5**). However, these networks were spatially distinguishable in both cerebral and cerebellar cortices in humans ^49^. One major factor may account for the discrepancy is the expansion of the prefrontal and parietal cortices in humans, resulting in more developed and well-separated networks for association brain areas, including default mode network, frontoparietal network, dorsal attention network and ventral attention network ^35^. In contrast, the prefrontal and parietal cortices of marmosets and other small non-human animals are relatively small, and these networks are not as clearly spatially distinguishable as in humans ^50,51^ by resting-state functional connectivity, which partly explained the discrepancy between marmosets and humans. However, despite similarities in connectivity patterns between the frontoparietal-like and default-mode-like networks to the cerebellar cortex, their relative connectivity strengths were different. By using a winner-take-all approach to map their topological relationship to the cerebellar cortex, we found results similar to those in humans: the frontoparietal-like and default-mode-like network occupied adjacent but distinct zones in non-motor areas of the cerebellar cortex (**Supplementary** Fig. S8). Thus, the two networks are spatially distinguishable the cerebellar mapping of marmosets, consistent with the results in humans ^49,52^.

The cerebello-cerebral gradient could not fully explain the intra-cerebral gradient patterns, which suggested that other anatomical features may exist to shape the functional connectivity of the cerebral cortex. As the cerebellar cortex has a remarkably uniform cytoarchitecture and local circuits, which anatomical features may contribute to the gradient patterns of the intra-cerebral connectivity? Recent advances in high-throughput transcriptional sequencing have started to unveil the molecular heterogeneity of the cerebellum ^53,54^. By single-cell RNA sequencing of different lobules across mouse cerebellum, a recent study achieved a comprehensive survey of cell types in the cerebellar cortex, and revealed considerable regional specialization in Purkinje neurons and gradient changes in gene expression for cerebellar interneurons ^54^. These molecular variations were associated with continuous changes in cell morphologies and electrophysiological properties. By spatially resolved transcriptomics, another study identified regional-specific gene expression patterns with the heterogeneity in Purkinje cells, which had not previously been described in single-cell sequencing studies ^55^. The molecular heterogeneity of the cerebellar cortex may be one of the anatomical features that contribute to the gradient patterns of functional connectivity ^56^. Future studies integrating both connectivity and gene expression data will be necessary to fully resolve the anatomical basis of the cerebellar functional gradients.

The cerebellar nuclei are key nodes for the cerebello-cerebral circuit loop, which receive inputs from the cerebellar cortex and send outputs to other brain regions ^24,36,57^. In the mammalian brain, there are three pairs of cerebellar nuclei, with the fastigial nuclei being considered as phylogenetically the oldest and the dentate nuclei the youngest ^36,58,59^. By brain-wide anterograde neuronal tracing, a previous study revealed distinct projection patterns of the three types of nuclei in mice, with their target regions being shifted relative to each other ^36^. Such shifts formed a gradient-changing pattern of connectivity to the cerebellar cortex, where fastigial (medial), interposed and dentate (lateral) nuclei innervated the vermis (motor), paravermis, and hemisphere (non-motor) of the cerebellar cortex, respectively.

Compared with mice brains, the primate cerebellar nuclei underwent nucleus-specific expansion during evolution (**Fig. 6C**). However, the connectivity of cerebellar nuclei has not been fully explored in the primate brains, for which the neuronal tracing approaches were less accessible. With the high-resolution diffusion MRI and awake resting-state fMRI data, we mapped topological connectivity patterns of the cerebellar nuclei in the cerebello-cerebral circuits of the marmoset brain (**Fig. 6**). The phylogenetically youngest dentate nucleus showed structural connectivity mainly to the Sim and Crus areas of the cerebellar cortex, which are non-motor areas at one extreme of the cerebello-cerebral functional gradient. In contrast, the phylogenetically oldest fastigial nucleus was mostly anatomically connected with the lobules I-VI, which are motor areas at the other extreme of the functional gradient. Consistently, the topological pattern was also reflected in the principal gradient of the functional connectivity between the cerebellar nuclei and the neocortex, with the dentate nucleus at one extreme and the fastigial nucleus at the other extreme (**Fig. 6F**). The connectivity patterns of the cerebellar nuclei of the marmoset revealed by diffusion tracking were similar to the patterns seen in the mouse as reported by neuro-tracing data ^36^, indicating the common topological principle of the cerebellar-nuclei connectivity across species. In all, these results provided a comprehensive map of anatomical and functional connectivity patterns of the marmoset cerebellum and characterized the close evolutionary relationship of the cerebello-cerebral circuits.

### Limitations and future directions

The Marmoset Brain Mapping V5 has provided the first comprehensive neuroimaging atlas of the marmoset cerebellum and revealed details of the cerebellar anatomy and connectivity. However, the V5 also faces limitations that may require attention and future modification. First, due to the resolution required to resolve the cerebellar anatomy, we chose to use the *ex-vivo* ultra-high resolution data of one brain sample to build the anatomical atlas rather than the *in-vivo* structural MRI data of a population ^12^. As the general neuroanatomy of the cerebellum is similar across animals, our atlas can be readily spatial registered to the data of other animals for most applications. However, it would be beneficial to create a population-based atlas based on *ex-vivo* ultra-high resolution data in order to reveal individual variations in the fine-detailed folia patterns. Second, based on awake resting-state fMRI, we were able to reconstruct the gradient patterns of the cerebellar functional connectivity. However, the functional domains of the cerebellum, including gross-level non-motor and motor presentations, were not defined and validated by multi-domain task fMRI data ^22^. Because of the practice difficulties in instructing animals to perform multi-domain tasks inside MRI scanners, we need other techniques, such as electrophysiology and spatial transcriptome, to explore the functional domains of the marmoset cerebellar cortex. Third, we estimated the connectivity between the cerebellar nuclei and the cerebellar cortex purely by diffusion tractography. Although our ultra-high resolution data improved the tracking accuracy and the cerebellar white matter is less complex than the cerebral cortex, diffusion tractography has inherent limitations and may not fully replicate real anatomical connections ^60^. It is important to examine the real connections underlying the topological relationship between the cerebellar cortex and nuclei by neuronal-tracing data and other cutting-edge techniques ^61–64^. Multi-omics integration of different types of data, including our current neuroimaging data, will be an important direction to fully characterize the anatomical and functional architecture of the cerebellum.

## STAR Methods

### Data Description

All marmoset data used in this study were collected from the *Marmoset Brain Mapping* project (*marmosetbrainmapping*.*org*), and all procedures were approved by the Animal Care and Use Committee (ACUC) of the Institute of Neuroscience, Chinese Academy of Sciences. The ultra-high resolution multi-modal *ex-vivo* data, from version 2 (MBMv2) of the *Marmoset Brain Mapping* project ^13^, were used to create the lobular parcellation, reconstruct fine-grained surfaces, and estimate structural connectivity. The *ex-vivo* data were collected on a 7T/30cm MRI (Bruker Biospin) with a 30-mm ID quadrature millipede coil (ExtendMR, LLC, Milpitas, CA), including 80 μm and 64 μm isotropic multi-shell diffusion MRI (dMRI), 80 μm isotropic MTR, and 50 μm isotropic T2*-weighted and T2-weighted images. The multi-shell dMRI had three b values and 204 DWI directions (8 b=0, 6 b=30, 64 b=2400, and 126 b=4800). A detailed scanning protocol was described in the MBMv2 resource paper ^13^. The awake resting-state fMRI data, from version 4 (MBMv4) of the *Marmoset Brain Mapping* project ^15^, were used to examine the functional connectivity and gradient patterns of the cerebellum. The data was collected from 39 marmosets, which were scanned repeatedly and resulted in a total of 710 fMRI runs (17 min per run). All resting-state fMRI data had a temporal resolution of TR = 2s, and a spatial resolution of 0.5 mm isotropic and were scanned using two opposite phase-encoding directions (left-to-right and right-to-left) to compensate for Echo Planar Imaging (EPI) distortions and signal dropouts. Besides, for each session, a T2-weighted image was scanned for spatial registration. A detailed scanning protocol of the resting-state fMRI can be found in the MBMv4 resource paper ^15^. Human data were from “The Hangzhou Normal University of the Consortium for Reliability and Reproducibility (CoRR-HNU) dataset” (Zuo et al. 2014), consisting of 30 young healthy adults (15 females, mean age = 24). The detailed information of the data can be found at http://fcon_1000.projects.nitrc.org/indi/CoRR/html/hnu_1.html.

### Lobular parcellations and surface reconstruction

To create a fine-detailed surface of the marmoset cerebellar cortex, the cerebellum was extracted from the 80 μm multi-modal templates of the MBMv2. The ultra-high resolution and multi-modalities of the MBMv2 provided rich contrasts to visually distinguish the fine-detailed folia patterns of the cerebellar cortex. According to the anatomical annotation of the Paxinos atlas ^19^ and folia patterns, the cerebellum was manually segmented into the gray matter and the white matter tissue types and parcellated into different lobules using ITK-Snap ^65^. The digital delineated lobules included I, II, III, IV, V, VI, VII, VIII, IX, X, SIM, Par, Cop, Fl, PFI, Crus I and Crus II, a total of 17 lobules. The I, II, III, and IV were merged into one region (I-IV), and the IX and X were merged into one (IX-X) to form a 13-lobule parcellation, similar to previous studies for the human cerebellar atlas. These lobule parcellations and tissue-type segmentation constituted the volumetric atlas of the marmoset cerebellar cortex.

Surfaces and flat maps were shown to be useful tools to visualize and analyze cerebellar imaging data ^31,32,67,68^. Based on the volumetric atlas, fine-detailed surfaces of the cerebellar cortex were generated using the Freesurfer ^30^. As tailored for *in-vivo* structural images of human brains, the Freesurfer would not work appropriately on our high-resolution *ex-vivo* images of the marmoset cerebellum. Therefore, related images were modified to conform to the requirements of the Freesurfer. First, the T2*-image, the gray matter, and the white matter segmentation maps of the cerebellum were resampled to a RSP orientation (Right-to-left, Superior-to-inferior, and Posterior-to-anterior), and the resolution information in image headers was modified from 80 μm x 80 μm x 80 μm to 1 mm x 1 mm x 1mm (note that only the header was modified, the images themselves were not downsampled). The value of the white matter segment was set to 127 or 255 for the surface tessellation (equivalent to the *mri_fill*) to create an original white matter surface of the cerebellum. Then, the gray matter and the white matter segments were merged, assigned specific values for each tissue type, and smoothed into a combined image. The pial surface of the cerebellar cortex was generated from the original white matter surface by the *mris_make_surfaces* command with modified parameters. The paired two surfaces were smoothed and manually adjusted to remove errors during surface generation. To create the flat map of the cerebellar cortex, the white matter surfaces were inflated by the *mris_inflate*, and sphere surfaces were created by the *mris_sphere*. Volumetric lobular parcellations were mapped to these surfaces by the ribbon method of the *wb_command* of the Connectome Workbench ^69^. Guided by the curvature map and the lobular parcellations, the inflated surface was manually cut and flattened by the *mris_flatten*. The raw flat map was then manually optimized using an in-house method to improve the visualization. Together, the white matter surface, pial surface, flat map, associated curvature map, and lobular-parcellation surface map, constituted the surface-based tool of our atlas.

Our atlas was generated from *ex-vivo* MRI data of brain samples, and the brain sample may expand or shrink during the fixation, preparation, and storage. When using the *ex-vivo* data to estimate the volume of the cerebellum, the expansion or shrinkage of the volume should be corrected. Thus, we calculated the total volumes of the cerebellum using the *ex-vivo* data of the MBMv2 template ^13^ and the population *in-vivo* templates of the MBMv3 ^12^, respectively. The volume ratio between the *ex-vivo* data and *in-vivo* data was then used to correct the expansion or shrinkage effect when estimating the volume of the cerebellar lobules.

All surfaces and flat maps were released in the standard GIFTI format, which is compatible with commonly-used surface-based visualization and analyzing tools, including Connectome Workbench, AFNI/SUMA ^70^, and Freesurfer ^30^. The lobular parcellation was also mapped onto the high-resolution surfaces of the cerebellar cortex and released in NIFTI format and GIFTI format.

### Functional connectivity and gradient analysis of the cerebellar cortex

Based on our recently published resting-state fMRI resource (MBMv4) of the marmoset ^15^, we analyzed the functional connectivity patterns of the marmoset cerebellum. The preprocessed data “regrBMWC0” of the MBMv4 were used in the following analysis, which represented data with band-passing and temporal denoising of motion parameters, motion-censors, white matter signals, and cerebrospinal fluid (CSF) signals. The above nuisance signal regression (motions, white matter signals, and CSF signals) and bandpassing filtering were conducted in a single regression model by the “3dDeconvolve” and “3dTproject” commands of AFNI. The preprocess of the human data was similar as the marmoset, including slice timing correction, motion correction, spatial normalization into MNI space, resampling to 3 × 3 × 3 mm voxels and smoothing with a Gaussian kernel (FWHM = 6 mm). Friston-24 parameters of head motion, white matter and ventricle signals were regressed out, followed by linear drift correction and temporal filtering (0.01 - 0.1 Hz).

The diffusion map embedding was used to estimate the functional-gradient patterns of the cerebellum ^33,34^. Similar to a previous study on gradients of the human cerebellum ^25^, we analyzed the functional gradients on two types of connectivity: 1) the intra-cerebellar functional connectivity that included only the vertices of the cerebellar cortex, emphasizing the intrinsic organization of the cerebellar cortex; and 2) the functional connectivity between the cerebellar cortex and the cerebral cortex, emphasizing the cerebello-cerebral communications.

To obtain the functional gradients, the preprocessed cerebellar data were first mapped to the surface of the cerebral cortex and smoothed with a 1mm FWHM kernel on the surface. To improve computational efficiency, 5000 regions of Interest (ROIs) were generated on the cerebellar inflated surface based on their spatial information (10000 and 20000 ROIs were also evaluated and the resulting functional gradients were similar to the 5000 ROIs). The Pearson correlation between each pair of cerebellar ROIs was calculated to form the matrix of the intra-cerebellar functional connectivity. The correlation between cerebellar ROIs and voxels of the cerebral cortex was calculated to form the matrix of cerebello-cerebral functional connectivity. The diffusion map embedding method was applied to these matrices by the BrainSpace ^34^, which nonlinearly dimensionality reduced the matrices into several orthogonalized gradients accounting for as much data variance as possible. The gradient analysis was performed on population-averaged correlation matrices and on the correlation matrices of each marmoset to examine the individual variabilities for each gradient. Take the principal gradient (gradient-1) for example, we calculated the mean and standard deviation (variance) value from the gradient-1 maps of all marmosets and then divided the variance by the mean to obtain a map that reflected the individual variabilities of the gradients.

To examine cerebellar functional connectivity to the 15 functional networks of the marmoset cerebral cortex from awake resting-state fMRI ^15^, we calculated the Pearson correlation between the time series of each cerebellar vertices and the mean time series of each contralateral network, which generated a 286308 (vertices) x 15 (networks) matrix. For each vertex, we calculated the mean and variance of the correlations to the fifteen networks and then divided the variance by the mean to obtain a map of the coefficient of variation. The map reflected the specificity of the functional connectivity between these networks and the cerebellar cortex.

### Structural-connectivity-based mapping and functional connectivity analysis of the cerebellar nuclei

The cerebellar nuclei were manually delineated on the MRI templates of the MBMv2 (80-μm isotropic resolution) ^10^, including the dentate nucleus, interposed nucleus, and fastigial nucleus, using the ITK-Snap ^65^. For cross-species comparisons, the three nuclei were also delineated on previously published *ex-vivo* MRI templates of C57BL/6 mice (30 μm isotropic resolution) ^71^ and *ex-vivo* MRI data of rhesus macaque (250 μm isotropic resolution) ^13^.

To map the topological relationship of the connectivity between cerebellar nuclei and the cerebellar cortex, diffusion tractography was used to estimate their structural connectivity based on ultra-high resolution dMRI data. The main results reported were based on the 80 μm dMRI data of the MBMv2 resource ^10^. To verify the results, we also performed a similar analysis on the 150 μm high-b-value dMRI data of the MBMv1 ^14^. The detailed data collection and preprocessing protocols were described in our previous paper ^66^.

The software Mrtrix3 was used for all diffusion tracking ^26^. Following its standard pipeline, the *dwi2response* was used to estimate the response function of the preprocessed dMRI data by the *dhollander* method. The *dwi2fod* was used to calculate the fiber orientation distributions (FOD) images by the spherical deconvolution of the multi-shell multi-tissue CSD algorithm. The FOD images were then used for diffusion tracking by the iFOD2 algorithm implemented in the *tckgen* command. Each voxel of nuclei was used as the seed and the tracking was performed within the cerebellum by the *tckgen* to obtain the tracking probability map of each nucleus voxel. For each probability map, the percentages of the selected streamlines that reached different cerebellar lobules were calculated. A winner-takes-all strategy was then applied to the maps, which assigned each nucleus voxel with the voxel of the cerebral cortex that had the largest tracking probability. As such, a topological map between the cerebellar cortex and the cerebellar nuclei was built by structural connectivity. Based on the topological correspondence, the lobule parcellation and functional gradients of the cerebellar cortex were projected to the cerebellar nuclei. To reveal the functional relationship between the cerebellar nuclei and the cerebral cortex, we calculated the functional gradient patterns of the two regions with the same method described previously ^25^. In brief, the fMRI data of the nuclei were smoothed with a 0.1 mm FWHM kernel (because of the small size of the nuclei, we used a small smooth kernel). The functional connectivity between each nucleus voxel and the voxels of the cerebral cortex was calculated. The diffusion map embedding method was then applied to the correlation matrices to obtain the functional gradients of the connectivity between the cerebellar nuclei and the cerebral cortex. We then examined the functional connectivity maps of the positive and the negative extremes of the principal gradient. The mean time series of the top 10% voxels of the two extremes were extracted respectively and their correlations to the cerebral cortex were calculated and converted with the Fisher Z-transformation. A paired two-sample t-test was used to compare the functional connectivity maps of the two extremes, and the multiple comparisons were corrected by the false discovery rate (FDR, adjusted P value < 0.05).

## Acknowledgments

We thank Dr. Ian Logan for the language editing and commenting on the manuscript. The study was supported by the grants from by the National Science and Technology Innovation 2030 Major Program (Grant No. STI2030-2021ZD0203900, 2022ZD0205000, and 2021ZD0200900), the National Natural Science Foundation of China (No. 32171088 to C.L.), the Lingang Laboratory Grant (No. LG-QS-202201-02 to C.L.), the Shanghai Municipal Science and Technology Major Project (No. 2018SHZDZX05 to C.L.),

## Author contributions

C.L. and X.Z. designed the study; X.Z., C.L., Y.Z., and C.W. analyzed the data; X.Z., C.L., and H.Y. constructed the MBMv5 atlas; F.F., C.L., and X.Z. designed the MBMv5 online atlas viewer; X.Z. and C.L. wrote the original manuscript; C.L., X.Z., Y.Y.G., and Y.Z. revised the manuscript; and C.L. and Y.Y.G. supervised the study.

